# APR-246 reactivates mutant p53 by targeting cysteines 124 and 277

**DOI:** 10.1101/214049

**Authors:** Qiang Zhang, Vladimir J.N. Bykov, Klas G. Wiman, Joanna Zawacka-Pankau

## Abstract

The *TP*53 tumor suppressor gene is frequently inactivated in human tumors by missense mutations in the DNA binding domain. *TP*53 mutations lead to protein unfolding, decreased thermostability and loss of DNA binding and transcription factor function. Pharmacological targeting of mutant p53 to restore its tumor suppressor function is a promising strategy for cancer therapy. The mutant p53 reactivating compound APR-246 (PRIMA-1^Met^) has been successfully tested in a phase I/IIa clinical trial. APR-246 is converted to the reactive electrophile methylene quinuclidinone (MQ), which binds covalently to p53 core domain. We identified cysteine 277 as a prime binding target for MQ in p53. Cys277 is also essential for MQ-mediated thermostabilization of wild-type, R175H and R273H mutant p53, while both Cys124 and Cys277 are required for APR-246-mediated functional restoration of R175H mutant p53 in living tumor cells. These findings may open opportunities for rational design of novel mutant p53-targeting compounds.

## Introduction

Tumor suppressor p53 is a transcription factor that acts as a sensor of multiple stress stimuli, e.g. DNA damage, hypoxia and oncogenic stress. Depending on the type and severity of the stress and other factors, p53 triggers distinct cellular responses including cell cycle arrest, senescence and apoptosis^1, 2^. More recent studies have shown that p53 also has roles in metabolism^3^, stem cell division^4^, fertility^5^ and cell death by ferroptosis^6^. The *TP*53 gene is inactivated by mutation in a large fraction of human tumors^7, 8^. The majority of *TP*53 mutations are missense mutations resulting in substitution of amino acid residues that make direct contact with DNA, such as R248W and R273H, or residues that are important for the structural integrity of the core domain, e.g. R175H and R249S. This leads to loss of specific DNA binding ^9^.

The high frequency of *TP*53 mutations in human tumors has stimulated efforts to develop therapeutic strategies for targeting mutant p53 in cancer. Several low-molecular-weight compounds have been reported to restore wild-type function to mutant p53 including PRIMA-1 and the PRIMA-1 analog APR-246 (PRIMA-1^Met^) ^10, 11, 12, 13, 14, 15, 16, 17^. APR-246 has been tested in a phase I/IIa clinical trial in patients with hematological malignancies or prostate cancer^18^. APR-246 is converted *in vitro* and *in vivo* to methylene quinuclidinone (MQ), a Michael acceptor that reacts with thiols in cysteines in the p53 core domain^19^. However, the mechanism by which APR-246/MQ reactivates mutant p53 is not fully understood.

Many mutant p53 proteins in human tumors are thermodynamically unstable at body temperature^20^. Studies of temperature-sensitive mutants suggest that stabilization of conformation is critical for regaining wild-type p53 activity^21, 22^. Thus, pharmacological stabilization of mutant p53 should allow its functional rescue and efficient elimination of tumor cells^23^.

Here we have examined the role of the Michael acceptor activity of MQ for thermostabilization of wild-type (wt) and mutant p53 core domains and refolding of R175H mutant p53 in living cells. We also show that Cys277 is essential for MQ-mediated thermostabilization of R175H and R273H mutant p53 core domains, and that both Cys124 and Cys277 are required for APR-246-mediated R175H mutant p53 reactivation in tumor cells.

## Results

### MQ binds to the p53 core domain via Michael addition

We have previously shown that conversion products of PRIMA-1 covalently bind to cysteine residues in the p53 core domain and reactivate mutant p53 (ref. 19), but whether MQ, the active conversion product of APR-246, itself binds to the p53 core domain has not been unequivocally demonstrated. To address this, we analyzed MQ-treated p53 core domains by Nanomate Linear Ion Trap Orbitrap (LTQ) hybrid mass spectrometry. Accurate masses of the wt and R273H and R175H mutant p53 core domains are shown in Figure 1a. We incubated wt and R273H mutant p53 core domains (20µM) with 50µM, 100µM or 200µM MQ at 21°C and assessed the degree of thiol modification by LTQ-MS. The deconvoluted mass spectra showed that 32% of the wt and R273H p53 core domain proteins were modified by one MQ molecule when incubated with lower concentrations of the compound (Figure 1b), and that all wt and R273H p53 protein molecules had two MQ adducts upon incubation with 200µM MQ (Figure 1d). We did not detect modification of all 10 cysteines in p53 core at this range of MQ concentrations.

**Figure 1.**
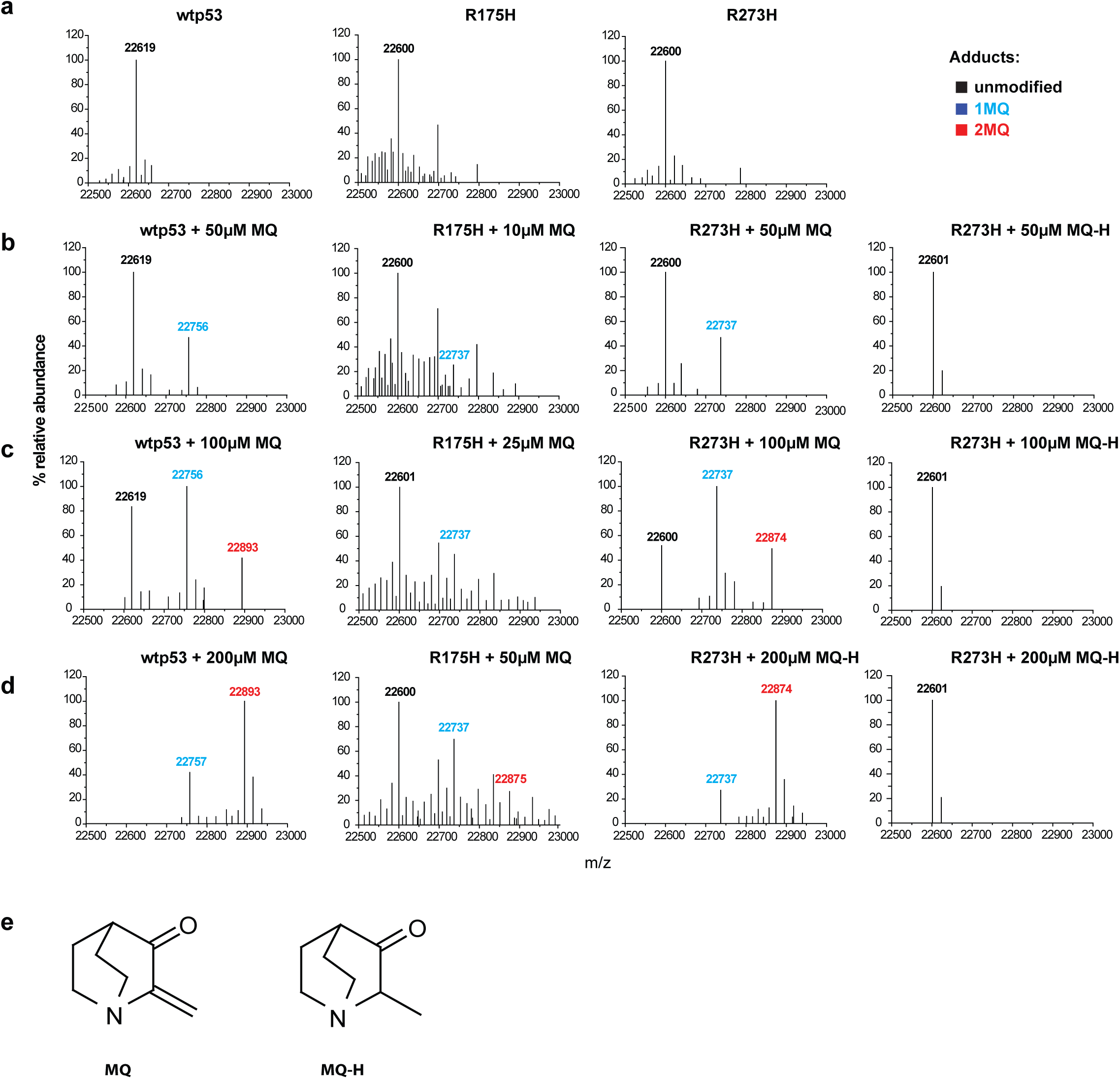
MQ binds to cysteine residues in wild-type and mutant p53 core domains in a dose-dependent manner. Mass measurement of wild-type, R273H and R175H p53 core domains by LTQ-MS. (a) mass spectra of p53 core domains. (b) – (d) reaction titration with MQ or MQ-H. p53 core domains were incubated with MQ at 50-200µM (wt and R273H) or 10-50µM (R175H) concentration ranges. One MQ adduct increased the molecular mass of p53 core domains by 137 Da. (e) structure of MQ and MQ-H.

Since we only obtained a low yield of the R175H p53 core domain, we analyzed it at lower protein concentration than wt and R273H (see Methods). Briefly, R175H core domain (3.2µM) was incubated with 10µM, 25µM or 50µM MQ. The deconvoluted mass spectra indicate that 20% of the R175H core was modified by one MQ molecule at 10µM (Figure 1c), and at 50µM MQ, 35% of the R175H core domain protein had one MQ adduct and 14% had two MQ adducts (Figure 1d). Thus, the number of p53 cysteine residues modified by MQ increased in a dose-dependent manner.

Next, we assessed binding of MQ-H, a hydrogenated analog of MQ that lacks the reactive carbon-carbon double bond (Figure 1e). We did not detect any modification at concentrations up to 200µM, indicating that MQ-H does not modify cysteine residues in p53. Thus, the Michael acceptor activity of MQ is required for modification of cysteines in p53.

### MQ enhances thermostability of the p53 core domain

To address the question if the modification of cysteine residues in p53 by MQ increases the thermostability of the p53 core domain, we applied differential scanning fluorimetry (DSF). This method allows analysis of the interactions between a protein and a ligand based on changes in the melting temperature of the protein (Tm). DSF demonstrated that the wt p53 core domain was the most stable of the three proteins with a Tm of 40.38±0.06ºC. Tm for the DNA contact mutant R273H was 39.16±0.52ºC, whereas the structural mutant R175H was the least stable protein with a Tm of 30.64±0.46ºC (Figure 2a). This is in a good accordance with previously published studies^20^. To further validate this result, we used circular dichroism (CD), which allows assessment of α-helix and β-sheet structures content. We performed CD analysis at 218 nm since both α-helix and β-sheet structures are detected at this wavelength. In agreement with our DSF data, CD measurements demonstrated that wt p53 core domain is the most stable protein followed by R273H core whereas the R175H core domain is considerably less stable. The Tm values are 44.74±0.08ºC, 43.96±0.37ºC and 36.24±0.33°C, respectively.

**Figure 2.**
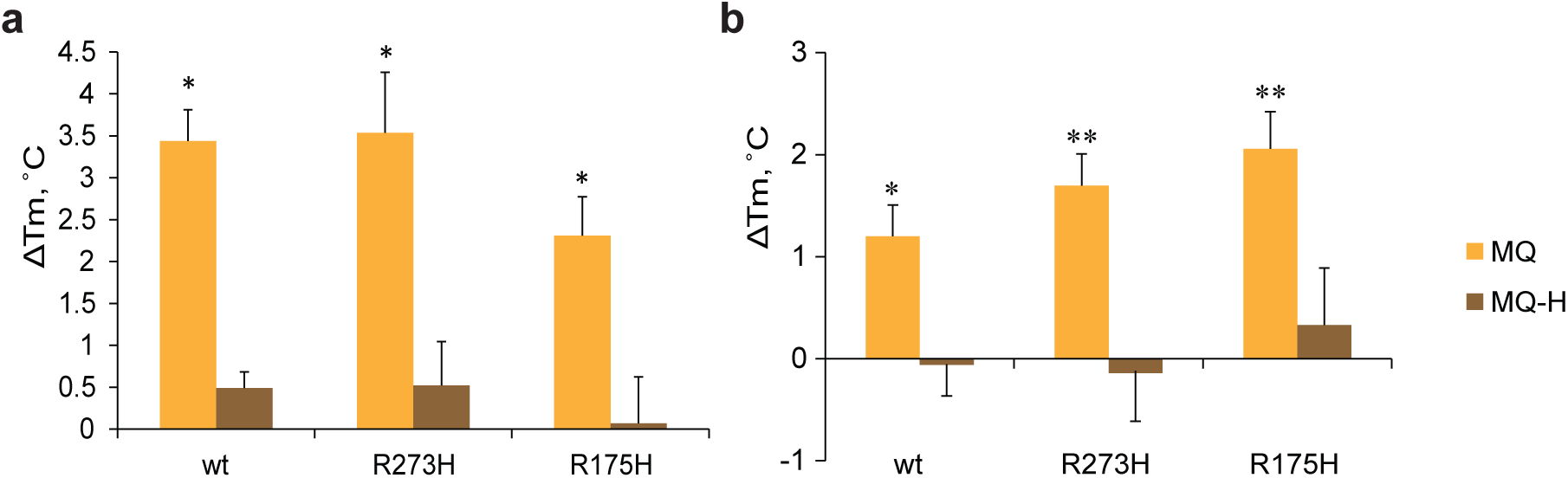
MQ modification of cysteine residues enhances p53 core domain thermostability. Changes in Tm after MQ or MQ-H modification were assessed by DSF (a) and CD at 218 nm (b) (mean±SD, n=3). All proteins were thermostabilized by MQ modification (yellow bars), but not by MQ-H (brown bars). * p<0.05, ** p<0.01 (student t-test).

We then incubated wt, R273H and R175H p53 core domains with 2mM MQ followed by DSF and CD analyses. This concentration of MQ was chosen to override the reducing agents DTT or TCEP that are included in the reaction buffer to maintain wild-type-like structure of the p53 core domains in solution. According to DSF, MQ increased the Tm values of wt, R273H and R175H p53 core domains by 3.44ºC, 3.54ºC and 2.31ºC, respectively (Figure 2a). This degree of p53 core domain thermal stabilization upon binding of a small molecule is in agreement with previously published results^16^. CD analysis confirmed thermostabilization of all three core domains by MQ as shown by the increase in Tm values by 1.20°C, 1.70°C and 2.06ºC for wt, R273H and R175H p53, respectively (Figure 2b). The inactive MQ analog, MQ-H did not significantly change the thermal stability of the p53 core domains as assessed by DSF (Figure 2a) and CD (Figure 2b).

Thus, MQ but not MQ-H increases the thermostability of the p53 core domain proteins, confirming that cysteine binding by Michael addition is critical for p53 thermostabilization by MQ. The results from DSF and CD were fully consistent; the p53 protein melting temperatures determined by the two methods correlated with each other (r=0.999, p=0.007). DSF was chosen for further studies.

### Identification of MQ binding sites in the p53 core domain

The p53 core domain has 10 cysteine residues with varying solvent accessibility. Previous studies have indicated that in the absence of DNA Cys277 has the highest solvent accessibility, followed by Cys182 and Cys229, whereas Cys135, Cys141 and Cys275 have poor solvent accessibility ^24^. Cys124 is located at the center of the flexible L1/S3 pocket, which can be stabilized by second-site mutations to rescue mutant p53 folding^25, 26, 27^. Interestingly, Cys124 shows a nuclear magnetic resonance (NMR) chemical shift upon binding of the CDB3 peptide that stabilizes mutant p53 (ref. 28). In addition, mutation at Cys124 was reported to abrogate reactivation of R175H mutant p53 by PRIMA-1 (ref. 25).

Thus, to investigate if the most solvent-exposed cysteine residues and Cys124 are critical for MQ binding to the p53 core, we first introduced single Cys to Ala substitutions at position 124, 182, 229 or 277 in the wt, R175H and R273H p53 core domains, and double substitutions at Cys124 and Cys277 in the same three core domain proteins. The R175H p53 core domain has low intrinsic thermostability and additional amino acid substitutions might destabilize it further. This probably explains why we only obtained negligible protein yield for the R175H-C124A and R175H-C124A-C277A core domains. Our LTQ-MS analysis of R273H mutant p53 core domains identified one MQ adduct in the R273H, R273H-C124A, R273H-C182A, and R273H-C229A p53 cores after incubation with 50µM MQ, whereas no MQ modification of the R273H-C277A and R273H-C124A-C277A core domains was detected (Figure 3a). However, incubation with 100-200µM MQ resulted in two to five MQ adducts in all p53 mutant core domains tested except R273H-C277A (Figure 3 b-c). This implies that MQ modifies several cysteines in p53, and suggests that Cys277 is the most reactive.

**Figure 3.**
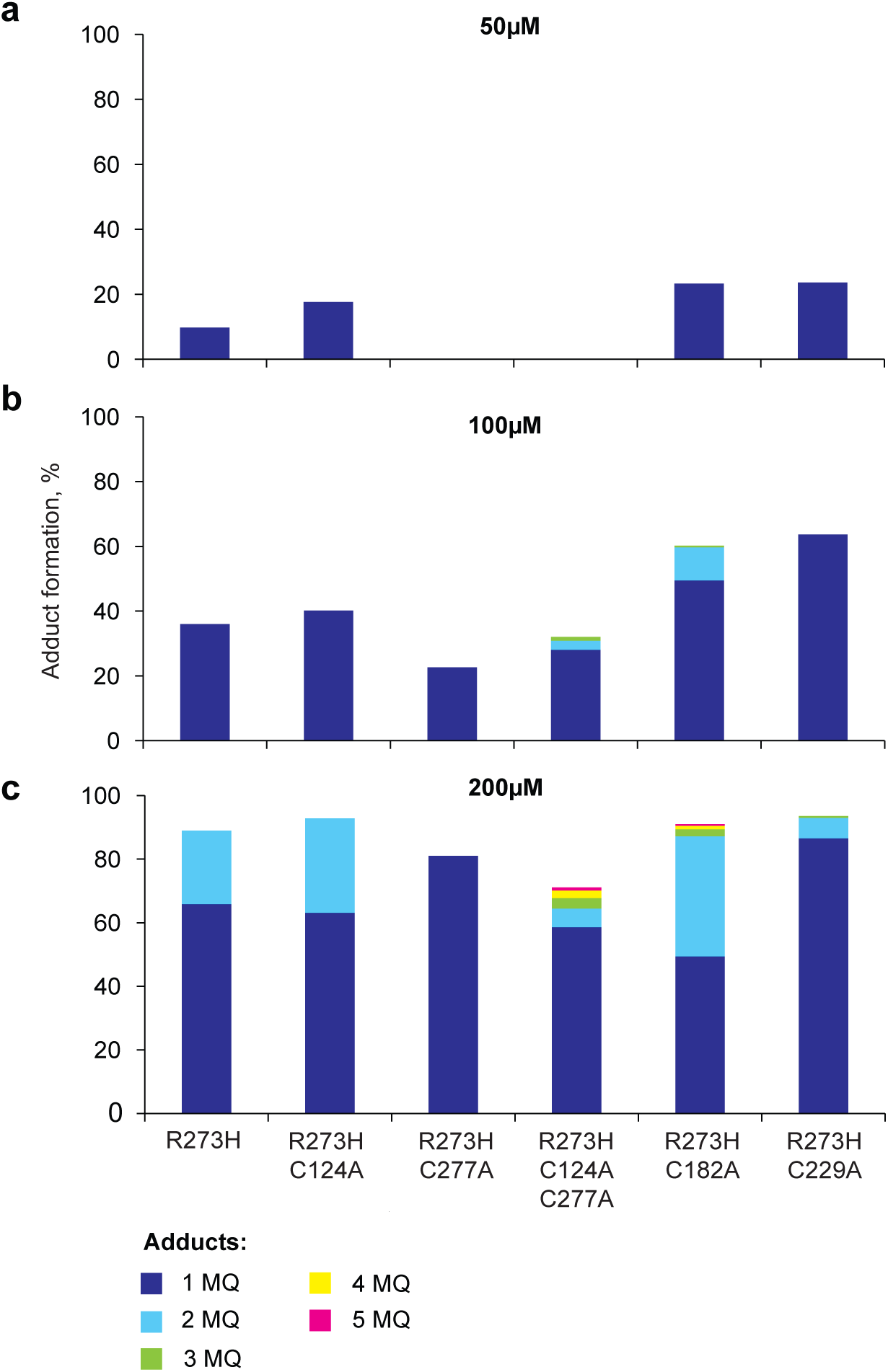
Cys277 is a prime binding site for MQ in the p53 core domain. Mass measurement of p53 core domains as assessed by LTQ-MS. Indicated p53 core domain proteins were incubated with MQ at increasing concentrations and mass shift was assessed. Proportion of MQ modified proteins was calculated based on relative intensity of each detected mass.

### Cys277 is essential for MQ-mediated thermostabilization of p53 core domains

In order to assess the role of selected Cys residues in MQ-mediated thermostabilization, we incubated wt, R175H and R273H p53 core domains carrying indicated Cys to Ala substitutions with MQ and performed DSF. The Tm values of the wt and p53 core domains carrying C124A, C277A, C182A, C229A and C124A-C277A substitutions were 42.36°C, 36.67°C, 43.07°C, 42.43°C, 40.59°C and 40.91°C, respectively (Table 1). MQ modification increased the Tm values of the C124A, C182A and C229A p53 core domain proteins by 5.82°C, 1.35°C and 1.00°C (Figure 4a), respectively. In contrast, MQ caused a slight destabilization of the C277A core domain by −0.06°C, and only stabilized the C124A-C277A core domain by 0.16°C, indicating that Cys277 is critical for MQ-mediated thermostabilization of p53.

**Table 1.** Melting temperatures (Tm) of p53 core domain proteins with indicated Cys to Ala substitutions as assessed by DSF.*

| **T<sub>m</sub>(°C)** | **wt** | **C124A** | **C277A** | **C182A** | **C229A** | **C124A-C277A** |
| --- | --- | --- | --- | --- | --- | --- |
| **wt** | 42.36±0.11 | 36.67±1.29 | 43.07±0.26 | 42.43±0.12 | 40.59±0.12 | 40.91±0.26 |
| **R175H** | 31.09±0.23 | - | 29.05±0.50 | 29.49±0.37 | 28.16±0.46 | - |
| **R273H** | 40.49±0.93 | 39.17±0.08 | 41.48±0.23 | 40.52±0.06 | 38.60±0.08 | 39.48±0.10 |

\*Sufficient amounts of the R175H-C124A and R175H-C124A-C277A mutant proteins could not be obtained for DSF analysis.
The differences in the T<sub>m</sub> values of wt, R175H and R273H p53 core domains between Table 1 and data presented in the main text (Results p. 5-6) are due to different batches of protein and iCycler instruments for DSF.

**Figure 4.**
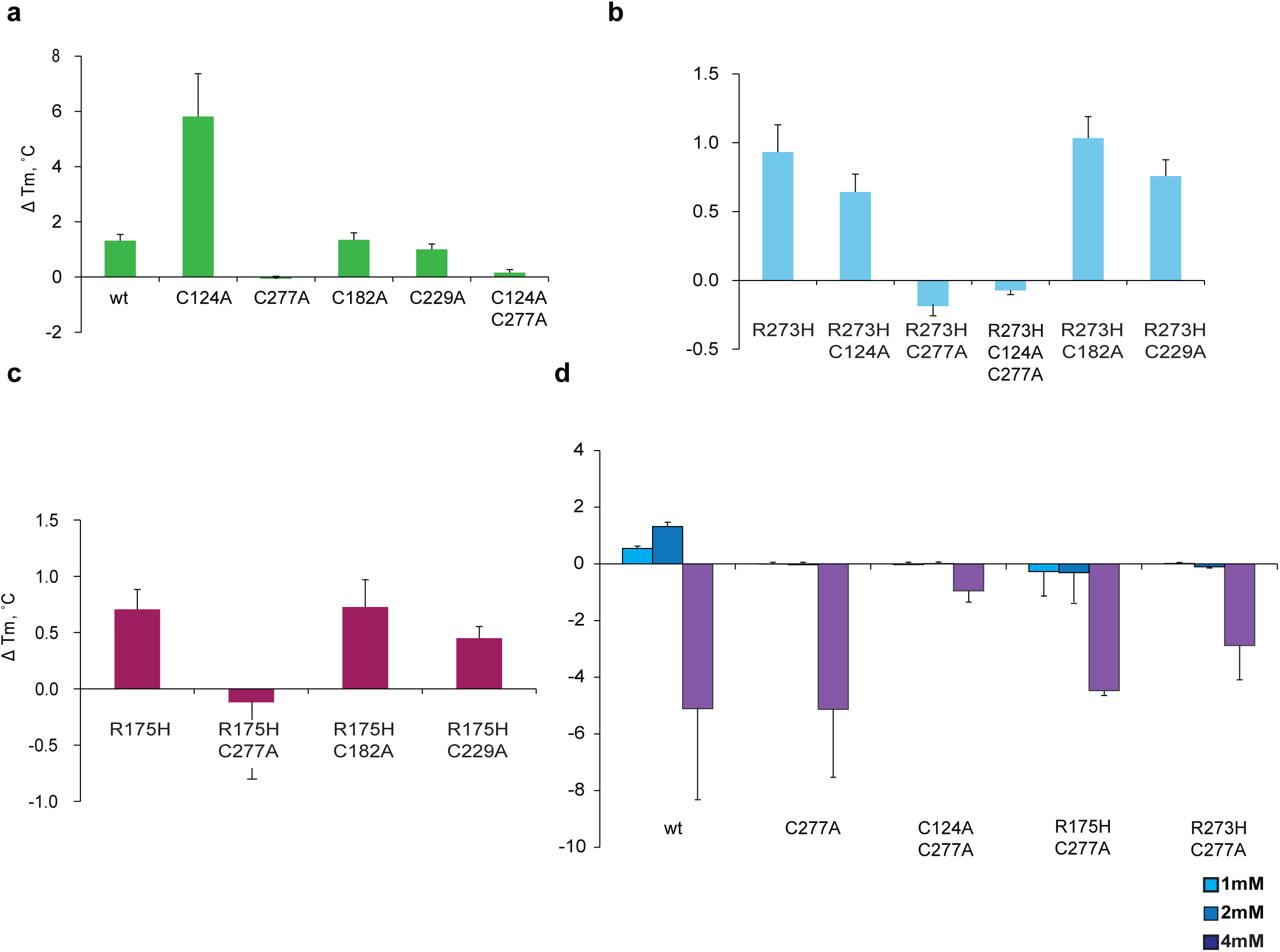
Cys277 is critical for MQ-mediated p53 thermostabilization. C277A substitution completely abolished MQ-mediated thermostabilization whereas other substitutions had little or no effect on wt (a), R273H (b) and R175H (c) core domains at 2 mM concentration. Higher concentrations of MQ (1, 2 or 4 mM) did not thermostabilize the indicated p53 core domain proteins with C277A substitution (d).

We then analyzed R273H p53 cores with the same C124A, C277A, C182A and C229A substitutions. The melting temperatures of the R273H, R273H-C124A, R273H-C227A, R273H-C182A and R273H-C229A p53 core domains were 40.49°C, 39.17°C, 41.48°C, 40.52°C and 38.60°C, respectively (Table 1). MQ treatment shifted the Tm values by 0.93°C, 0.64°C, −0.19°C, 1.03°C and 0.76°C (Figure 4b), respectively. The Tm of the R273H core domain with C124A-C277A double substitution was 39.48°C and only changed by −0.07°C upon MQ modification.

As indicated above, we were not able to obtain sufficient amounts of the R175H-C124A and R175H-C124A-C277A p53 core domains. Thus, only the R175H, R175H-C277A, R175H-C182A and R175H-C229A core domains were further analyzed. Their respective melting temperatures were 31.09°C, 29.05°C, 29.49°C and 28.16°C (Table 1). MQ modification shifted the melting temperatures by 0.71°C, −0.12°C, 0.73°C and 0.45°C (Figure 4c), respectively. Thus, C227A substitution abrogated MQ-induced thermostabilization in wt as well as R273H and R175H mutant p53 core domain backgrounds.

Our MS data demonstrate that MQ binds to the p53 core domain in a concentration-dependent manner and that high concentrations of MQ lead to modification of more than one cysteine residue in p53. This raises the question whether high concentrations of MQ might induce p53 thermostabilization even in the absence of Cys277. To address this, we incubated wt, C277A, C124A-C277A, R175H-C277A and R273H-C277A p53 core domains with 1 mM, 2 mM or 4 mM MQ and assessed protein thermostability by DSF (Figure 4d). The highest concentration of MQ (4 mM) destabilized all core domains (Figure 4d). The wt p53 core domain was stabilized by MQ at 1 and 2 mM, as observed previously. However, p53 core domains with the C277A substitution were not stabilized at the same concentrations. Thus, this result further supports our conclusion that Cys277 is indispensable for MQ-mediated p53 core domain stabilization.

### APR-246 and MQ induce R175H mutant p53 refolding to wild-type conformation

Proper folding of p53 is crucial for sequence-specific DNA binding and transactivation of p53 target genes. As a rule, *TP*53 mutations in tumors lead to p53 unfolding and loss of sequence-specific DNA binding capacity. PRIMA-1 was shown to promote R175H mutant p53 refolding to wild-type conformation in SKOV-His175 cells^11^. However, this has yet to be demonstrated for APR-246 and MQ.

The wild-type p53 conformation-specific monoclonal antibody PAb1620 allows detection of mutant p53 refolding to wild-type-like conformation. We verified specificity of the PAb1620 antibody in immunostaining by treating HCT116 human colon carcinoma cells with doxorubicin to induce the levels of wild-type p53. HCT116 cells expressing wild-type p53 or Saos-2 cells expressing R273H mutant p53, which retains wild-type-like conformation, showed PAb1620+ staining (Supplementary Figure 1). In contrast, H1299 cells expressing R175H mutant p53 were PAb1620 negative. Treatment with APR-246 induced positive PAb1620 staining in H1299-R175H cells, suggesting that APR-246 via MQ restores wild-type conformation of this mutant (Supplementary Figure 1).

To determine if APR-246 and MQ can refold endogenous R175H mutant p53 in TOV-112D ovarian carcinoma cells, we treated the cells with APR-246 or MQ and performed co-immunostaining with PAb1620 and anti-p53 polyclonal antibody FL-393. In parallel, the cells were stained with the monoclonal HO3.5 antibody that specifically detects unfolded p53 conformation in a manner similar to PAb240 (ref. 29). APR-246 treatment increased PAb1620 staining (Figure 5a) and decreased HO3.5 staining (Figure 5b), indicating refolding of R175H mutant p53 to wild-type-like conformation.

**Figure 5.**
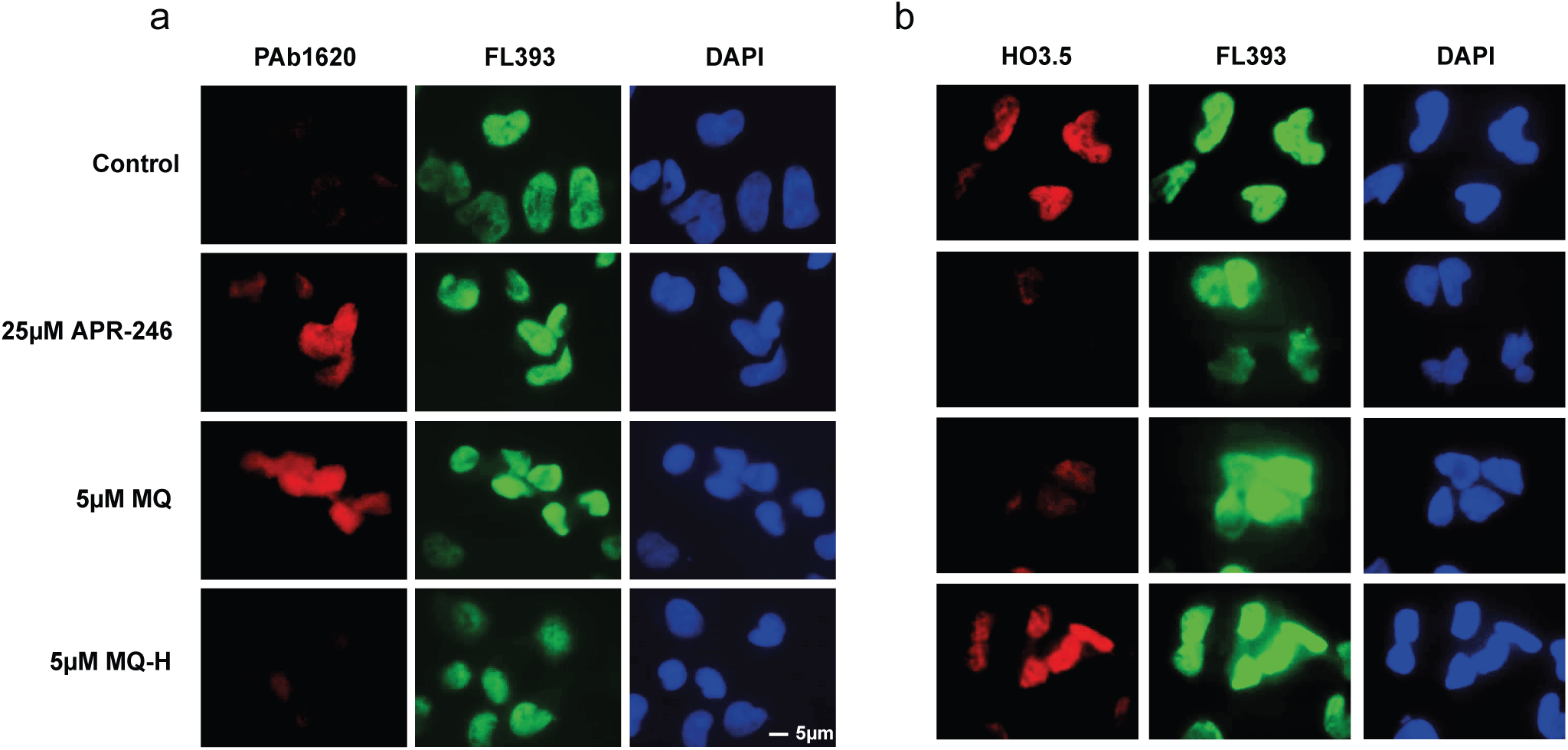
APR-246 and MQ enhance wild-type p53 conformation-specific PAb1620 epitope in tumor cells carrying R175H mutant p53. (a) Immunofluorescence staining of TOV-112D cells treated with APR-246, MQ or MQ-H using wild-type p53 conformation-specific antibody PAb1620 and co-immunostaining with general p53 antibody FL-393. (b) Immunofluorescence staining of TOV-112D cells treated with APR-246, MQ or MQ-H using the mutant p53 conformation-specific antibody HO3.5 and co-immunostaining with general p53 antibody FL-393.

Next, we treated TOV-112D cells with MQ and stained with PAb1620 and HO3.5 antibodies. Like APR-246, MQ induced PAb1620 staining (Figure 5a), which coincided with decreased HO3.5 staining (Figure 5b). We did not detect any changes in PAb1620 and HO3.5 staining after treatment with MQ-H, confirming that the Michael acceptor activity of MQ is crucial for mutant p53 refolding in living tumor cells.

### Cysteines 124 and 277 are important for APR-246/MQ-mediated R175H mutant p53 reactivation

To investigate the role of cysteine residues in mutant p53 reactivation by APR-246/MQ in tumor cells, we transiently transfected p53 null H1299 cells with vectors encoding R175H, R175H-C124A, R175H-C277A or R175H-C124A-C277A p53 mutant proteins, or with control vector (pCMV). Western blotting confirmed similar levels of expression of p53 in all transfectants (Supplementary Figure 2). Next, the cells were treated with 45µM APR-246. We chose this concentration since it induced cell death in R175H mutant p53-transfected cells but only marginally affected empty vector-transfected cells. We then assessed apoptosis and expression of p53 targets p21, Fas and Bax by flow cytometry. Signals obtained in mock-treated control cells were subtracted from the corresponding signals obtained in APR-246-treated cells, and the values are presented as relative increase, namely ΔAnnexin V, Δp21, ΔFas and ΔBax. Cells transfected with R175H showed substantial induction (by 61.06%) of Annexin V staining after APR-246 treatment when compared to cells transfected with pCMV empty vector (3.91%). We observed relatively low induction of Annexin V staining in cells transfected with R175H-C124A p53 (13.96%), whereas cells expressing R175H-C277A or R175H-C124A-C277A p53 showed no induction of Annexin V staining as compared to control vector-transfected cells (Figure 6a). The p53 targets p21 (Figure 6b) and Bax (Figure 6c) were highly induced by APR-246 in cells expressing R175H p53 but only slightly or moderately increased in cells expressing the R175H-C124A and R175H-C124A-C277A mutants. Cells expressing R175H-C277A mutant p53 showed no induction of p21 or Bax. The p53 target Fas (Figure 6d) was strongly induced by APR-246 in the R175H-transfected cells but no induction compared to control vector-transfected cells was detected in cells expressing R175H-C124A, R175H-C277A or R175H-C124A-C277A p53 proteins, consistent with the observed absence of apoptosis induction.

**Figure 6.**
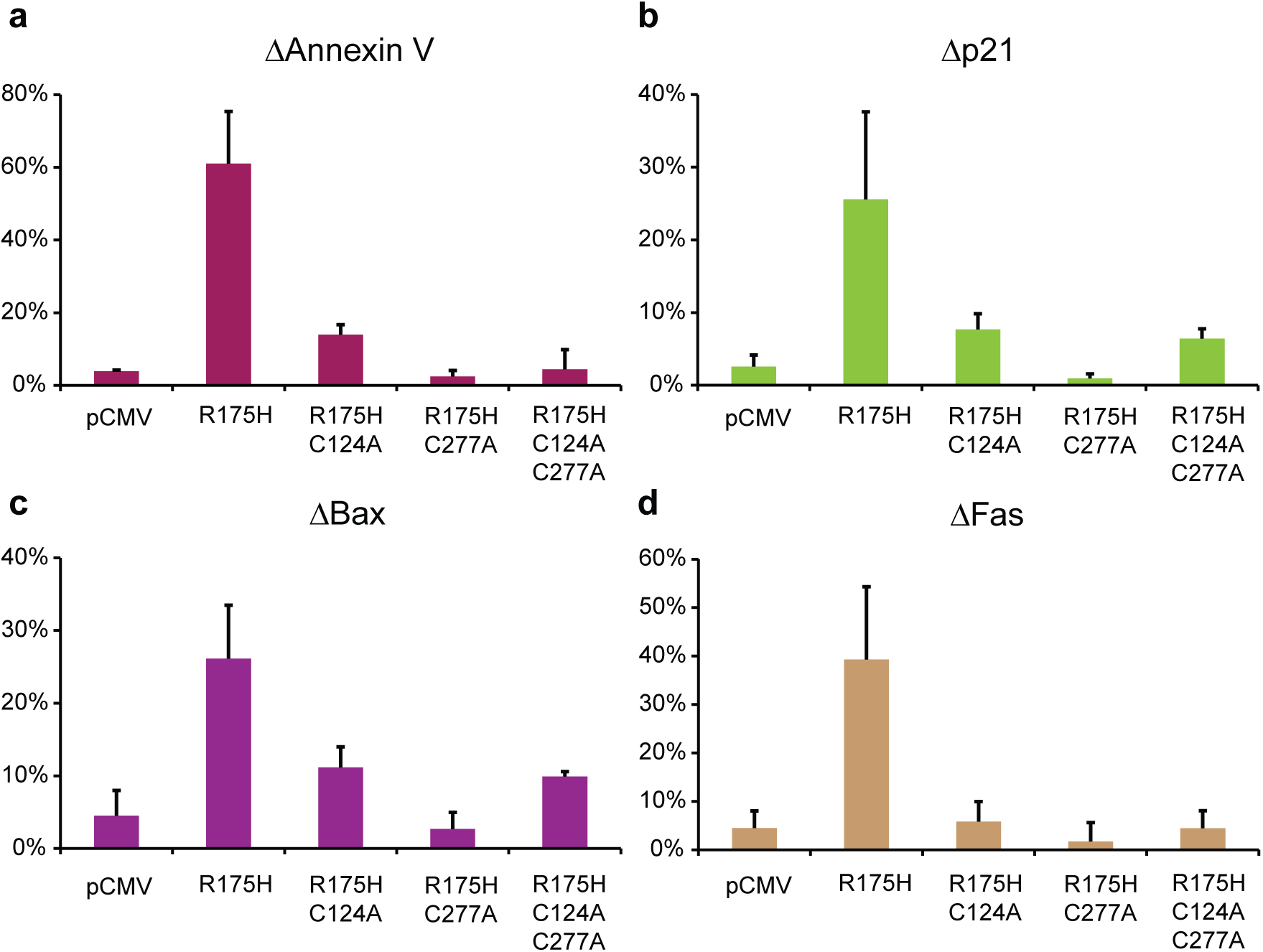
Cys124 and Cys277 are crucial for APR-246/MQ-mediated R175H reactivation in living cells. H1299 cells expressing corresponding p53 mutant proteins were stained with Annexin V, p21, Fas and Bax and examined by flow cytometry. Both C124A and C277A abolished APR-246-induced apoptosis (a), and upregulation of p53 targets p21 (b), Bax (c) and Fas (d).

To determine whether the C124A and C277A substitutions themselves affect wild-type p53 function, H1299 cells transfected with wt, C124A, C277A, R175H and R273H p53 constructs were assessed for apoptosis and expression of p53 target gene p21 by flow cytometry. pCMV vector was used as a control. Induction of Annexin V (Supplementary Figure 3a) was detected in cells expressing wt, C124A or C277A p53 proteins, which coincided with slight induction of p21 (Supplementary Figure 3b), but not in cells transfected with R175H or R273H p53 constructs. Thus, C124A or C277A substitution *per se* does not inactivate p53 in this experimental system, supporting our conclusion that these cysteines play a key role in APR-246/MQ-mediated mutant p53 reactivation.

## Discussion

*TP*53 gene mutation occurs in around 50% of human tumors and is emerging as predictive biomarker for currently available cancer therapy. A number of *in vitro*, *ex vivo* and *in silico* approaches have been applied to identify small molecules that reactivate mutant p53 by restoring wild-type p53 conformation^10, 30^. The mutant p53-targeting compound APR-246 has been tested in a phase I/IIa clinical trial in patients with hematological malignancies or prostate cancer^18, 31^, and is currently being tested in a phase II clinical trial in patients with high-grade serous (HGS) ovarian cancer (see clincaltrials.gov). Both PRIMA-1 and APR-246 are converted to methylene quinuclidinone (MQ)^19^. MQ is a potent electrophile that has a highly reactive carbon-carbon double bond coupled to the electron-withdrawing carbonyl group, defining it as a Michael acceptor. At physiological conditions, thiols in proteins are good nucleophiles and therefore prime targets for electrophilic attack by MQ. Our earlier studies clearly demonstrated thiol modifications in the p53 core domain by PRIMA-1 conversion products^19^. This led us to conclude that APR-246-mediated mutant p53 reactivation involves covalent binding of MQ to p53. Several other mutant p53-reactivating compounds, such as MIRA-1 (ref. 32), CP-31398 and STIMA-1 (ref. 33), 3-benzoylacrylic acid^14^ and 2-sulfonylpyrimidines^16^, possess similar reactivity with thiols, indicating that the observed association between thiol reactivity and mutant p53 reactivation is not coincidental.

A few methods have been successfully used to measure protein thiol modifications including antibody-based detection or chemical tagging of the modified thiol groups^34^. However, the development of mass spectrometry (MS) methods allowing the analysis of proteins using electrospray ionization (ESI) coupled with high-resolution instrumentation has significantly advanced studies of thiol modifications. Here we applied LTQ-MS to assess p53 core domain thiol modifications. We show that the MQ analog MQ-H that lacks a reactive carbon-carbon double bond and therefore lacks Michael acceptor activity, unlike MQ itself, does not modify cysteine residues in the p53 core domain, does not enhance p53 thermostability and does not induce R175H mutant p53 refolding according to PAb1620 staining. Thus, by using several approaches, we demonstrate that the electrophilic properties of MQ are essential for cysteine modification, thermostabilization and refolding of mutant p53.

Although previous studies have indicated that PRIMA-1 conversion products bind covalently to the p53 core domain^19^, the exact cysteine target residues for the major PRIMA-1 and APR-246 conversion product MQ have remained unknown. We applied LTQ-MS analysis to a set of Cys to Ala mutants to identify cysteine residues that are critical for MQ binding and MQ-mediated stabilization of mutant p53. The reactivity of cysteine residues in a protein is largely affected by their solvent accessibility. Among 10 cysteines in p53 core domain, Cys176, Cys238 and Cys242 coordinate a zinc ion which is responsible for holding p53 loops together^9^, making them less likely targets for modification. Cys135, Cys141 and Cys275 are poorly accessible to solvent based on the X-ray crystal structure of the p53 core domain. Cys277 and Cys182 have the highest solvent accessibility, followed by Cys229 (ref. 24). Interestingly, Cys277 has the lowest pKa of all p53 cysteines, making it the strongest nucleophile in the protein. Thus, Cys 277 combines the greatest solvent accessibility with the highest nucleophilicity, suggesting that it might be a prime target for MQ.

Indeed, we found that Cys277 to Ala substitution abolishes MQ binding to p53 core domain, at lower concentrations. Moreover, the ability of MQ to thermostabilize p53 core domains is impaired in Cys277 to Ala p53 mutants. Thus, a good correlation exists between the extent of MQ adduct formation and MQ-mediated thermostabilization of the p53 core domain.

Cys182 and Cys277 have recently been identified as the prime binding sites for PK11000, a 2-sulfonylpyrimidine compound that reacts with cysteines and thermostabilizes p53 (ref. 16). It is noteworthy that although Cys277 interacts directly with DNA, modification of this residue by PK11000 did not change p53 DNA binding and transactivation of target genes. Our results demonstrate that substitution of Cys182 to Ala does not affect p53 modification and thermostabilization by MQ in any significant way, implying that Cys182 is not essential for mutant p53 reactivation by APR-246/MQ.

Kaar and colleagues^14^ identified 3-benzoylacrylic acid as a thiol-binding compound that reacts first with Cys124 and Cys141 and to a lesser extent with Cys135, Cys182 and Cys277 in p53. Cys124 was also identified as a target for PRIMA-1 by molecular modelling, and Cys124 to Ala substitution abolished PRIMA-1-induced reactivation of mutant p53 in human tumor cells^25^. Here, we examined the role of Cys124 and found that substitution of this cysteine did not impair MQ binding and p53 thermostabilization. However, Cys124 to Ala substitution abrogated R175H reactivation by APR-246/MQ in tumor cells, in agreement with the results of Wassmann et al.^25^.

In order to exclude the possibility that the Cys to Ala substitutions themselves impair wild-type p53 function in our experimental setting, we examined whether the Cys to Ala substitutions impair ability of p53 to transactivate p21 and induce apoptosis. We confirmed that these substitutions do not affect normal p53 function to any major extent, supporting the notion that the observed effects are indeed due to an important role of Cys124 and Cys277 for APR-246/MQ-mediated mutant p53 reactivation.

In conclusion, our data demonstrate that specific cysteines are critical targets for mutant p53 reactivation by APR-246/MQ. Our findings may open opportunities for designing novel compounds targeting mutant p53 based on a similar mechanism of nucleophilic addition at the identified binding sites.

## Materials and Methods

### Cell lines and reagents

Human lung adenocarcinoma cells H1299 and osteosarcoma cells Saos-2 are p53 null. The sub-lines H1299-R175H and Saos-2-R273H stably express the indicated mutants^11, 32^. Human HCT116 colon carcinoma cells express wild-type p53. Human epithelial ovarian cancer cells TOV-112D express R175H mutant p53. All cells were cultured at 37°C, 5% CO_2_ in IMDM medium (Hyclone, Logan, Utah) supplemented with 10% FBS (Thermo Fisher Scientific, Waltham, MA).

APR-246, MQ and MQ-H were obtained from Aprea Therapeutics AB, Stockholm, Sweden. Methanol, formaldehyde and acetonitrile were purchased from Thermo Fisher Scientific (Waltham, MA). Formic acid was purchased from Sigma-Aldrich (St. Louis, MO). Lipofectamine 2000 was from Thermo Fisher Scientific (Waltham, MA). All solvents were of analytical grade and are commercially available.

Rabbit polyclonal anti-p53 FL-393, rabbit polyclonal anti-GAPDH, mouse monoclonal anti-p53 DO-1 and mouse monoclonal PAb1620, Alexa Fluor 647 conjugated FL-393, Alexa Fluor 488 conjugated anti-p21 antibodies were from Santa Cruz Biotechnology (Heidelberg, Germany). Mouse monoclonal antibody HO3.5 was a gift from Professor Thierry Soussi, Karolinska Institutet. Polyclonal rabbit anti-Bax Biotin OAAF02999 and Qdot^TM^ 605 streptavidin were from Nordic Biosite (Stockholm, Sweden). BV510 mouse anti-human CD95 (Fas) and BD Horizon V450 Annexin V were from BD Biosciences (Stockhlom, Sweden).

### Site-directed mutagenesis

Prokaryotic and eukaryotic plasmid constructs were produced by Genscript, Piscataway, NJ.

### Expression and purification of proteins

p53 cores (94-292) were cloned into pNIC28-Bsa4 that adds an N-terminal hexahistidine tag and transformed into *E. coli* strain Rosetta2 (DE3). Bacteria were grown in TB medium supplemented with 8 g/l glycerol at 37ºC with shaking. Protein expression was induced with 0.5mM IPTG at 18ºC overnight. Afterwards bacteria were pelleted by centrifugation and lyzed in cold IMAC lysis buffer (50 mM TRIS, 300 mM NaCl, 10% glycerol, 0.05 mM ZnCl, 0.5 mM TCEP, pH 8.0) supplemented with complete protease mix (complete EDTA-free (protease inhibitor) and 5 μl benzonase nuclease (250 U) and stored at −80ºC. After thawing, the cells were lyzed by pulsed sonication (4s/4s 3 min, 80% amplitude), centrifuged (20 min at 49000 xg) and the soluble fractions were decanted and filtered through 0.45μm filters. The samples were loaded onto the ÄKTA Xpress LC and purified overnight. His-tag was cleaved with Thrombin. Sample homogeneity was confirmed by mass spectrometry and the concentration was measured by nanodrop. The proteins were aliquoted and stored at −80˚C in storage buffer (50 mM TRIS, 800 mM NaCl, 10% glycerol, 2.0 mM TCEP, pH 8.0).

### Mass spectrometry

Wild-type and R273H p53 core domains were de-salted against 20 mM ammonium acetate buffer by using 10K concentration columns (Vivaspin, GE Healthacare, Chicago, IL). Twenty µM of the purified protein were incubated with 0 µM (control), 50 µM, 100µM or 200µM MQ for 15min at.21°C. R175H core domains were de-salted by ZipTip C4 resin tips for MALDI-ToF MS (Merck Millipore, Billerica, MA) following the manufacturer protocol. 3.2µM of R175H protein were treated with 0µM (control), 10µM, 25µM or 50µM of MQ for 15min at 21°C. 5% formic acid (1:1 volume ratio) was added to the samples to increase the ionization sensitivity. Samples were analyzed by LTQ XL mass spectrometry (Thermo Fisher Scientific, Waltham, MA) fitted with an automated nanospray source (TriVersa Nanomate, Advion Biosciences, Ithaca, NY) using nanoelectrospray chips with spraying nozzels. The ion source was controlled using the Chipsoft 8.3.1 software (Advion Biosciences, Ithaca NY). Three microliters of each sample were loaded into a 96-well plate and injection volume was one and a half microliters. Full scan spectra were collected at the *m/z* 500–2,000 in positive ion mode. The mass spectra of each sample were acquired in profile mode over 4 min. The spectra were analyzed using XCalibur^TM^ Software (Thermo Fisher Scientific, Waltham, MA). Deconvoluted ESI spectra are presented.

### Circular dichroism

75 μg of p53 core domain proteins were incubated with or without 2 mM MQ in 250µl 40mM potassium phosphate buffer (pH 7.5) and 1 mM DTT for 1hr at 21ºC. CD measurements were performed on Jasco-810 (Jasco Inc., Tokyo, Japan) with 0.01 pathlength. Denaturation curves were obtained by measuring the circular dichroism spectra at 218 nm. Melting temperatures were analyzed by GraphPad Prism 6 (Graphpad Software Inc, La Jolla, CA) according to the Boltzmann equation 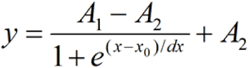, x_o_-inflection point.

### Differential scanning fluorimetry

5μg p53 core domains were incubated with 1 mM to 4 mM of MQ in 25μl 40mM potassium phosphate buffer (PH 7.5) with 1mM DTT for 1 hr at 21ºC under controlled conditions. 1 μl of 25x Sypro orange were added to each well. The fluorescence was assessed by Bio-Rad iCycler (Bio-Rad Laboratories, CA) at increasing temperature from 10ºC to 75ºC with a rate of 1ºC per min. Tm values were calculated by GraphPad Prism 6 (Graphpad Software Inc, La Jolla, CA).

### Immunofluorescence staining

Cells were plated into a 16 well chamber slide at a density of 3000 cells per well, allowed to attach overnight, and treated with 25µM APR-246 or 5µM MQ/MQ-H for 16hr. Cells were washed, fixed with 4% formaldehyde and permealized with 0.2% Triton X. Mouse PAb1620 or HO3.5 antibody were co-incubated with rabbit FL-393 antibody, all were diluted 1:200 in 2% BSA for 1hr at 4˚C. Anti-rabbit Alexa 488 and anti-mouse Alexa 594 conjugates were used as secondary antibody with 1:200 dilution in 2% BSA.

### Flow cytometry

Cells were grown on 6-well plates at an initial density of 500,000 cells/well. Sixteen hours later cells were transfected for 24 hr with p53 expression vectors or empty vector using Lipofectamine 2000 according to the manufacturer's protocol (Life Technology, Waltham, MA). The medium was then replaced with fresh medium, the cells were reseeded at a density of 20,000 cells/well after 6hr culture, and treated with APR-246 on the following day. Cells were collected 24 hr post-treatment, stained with Annexin V, fixed with 4% formaldehyde, permealized with 90% methanol and stained with Fas, Bax and p21 antibodies. Cells were analyzed on a NovoCyte Flow Cytometer (ACEA Biosciences, Solna, Sweden).

## Acknowledgements

This work was supported by grants from the Swedish Cancer Fund (Cancerfonden), the Swedish Medical Research Council (VR), the Cancer Society of Stockholm (Radiumhemmets Forskningsfonder), Åke Wiberg Stiftelse, The Strategic Research Programme in Cancer (StratCan), and Karolinska Institutet. We thank Prof. Thierry Soussi, Department of Oncology-Pathology, Karolinska Institutet, for generous gift of the HO3.5 antibody. We also thank the Protein Science Facility (PSF) and Proteomics Karolinska at the Department of Medical Biochemistry and Biophysics, Karolinska Institutet, for valuable help with p53 core domain purification and technical support for mass spectrometry.

